# The protease interpain A of *Prevotella intermedia* promotes human oral squamous cell carcinoma cells proliferation and migration

**DOI:** 10.1101/2024.10.20.619282

**Authors:** Szu-Min Chang, I-Hsiu Huang, Yun-Jhen Ciou, Wen-Rong Wang, Ni-Hung Wu, Chih-Pin Chuu, Jenn-Ren Hsiao, Jeffrey S Chang, Jang-Yang Chang, Jenn-Wei Chen

**Affiliations:** Institute of Basic Medical Sciences, College of Medicine, National Cheng Kung University, Tainan 70101, Taiwan; Department of Biochemistry and Microbiology, Oklahoma State University Center for Health Sciences, Tulsa, OK 74107, USA; Department of Microbiology and Immunology, College of Medicine, National Cheng Kung University, Tainan 70101, Taiwan; Institute of Cellular and System Medicine, National Health Research Institutes, Miaoli 35053, Taiwan; Department of Otolaryngology, National Cheng Kung University Hospital, Tainan 70101, Taiwan; National Institute of Cancer Research, National Health Research Institutes, Tainan 70101, Taiwan; Institute of Biotechnology and Pharmaceutical Research, National Health Research Institutes, Miaoli 35053, Taiwan; TMU Research Center of Cancer Translational Medicine, Taipei Medical University, Taipei 110301, Taiwan; Center of Infectious Disease and Signaling Research, National Cheng Kung University, Tainan 70101, Taiwan

**Keywords:** *Prevotella intermedia*, OSCC, interpain A, PAR-2

## Abstract

*Prevotella intermedia* has been increasingly recognized as a potential contributor to oral squamous cell carcinoma (OSCC), yet the underlying mechanisms remain poorly defined. In this study, we identified interpain A (InpA), a cysteine protease secreted by *P. intermedia*, as a key virulence factor that promotes oral epithelial cell proliferation and OSCC cell migration. Conditioned medium (CM) derived from *P. intermedia* strain ATCC 25611 enhanced proliferation in both normal oral keratinocytes (SG cells) and OSCC cell lines (SCC-15, SAS). The pro-proliferative effect was abolished upon heat inactivation of the CM and inhibited by E64, a cysteine protease inhibitor, and FSLLRY-NH_2_, a PAR-2 antagonist, suggesting the involvement of protease-activated receptor-2 (PAR-2) signaling. InpA was highly secreted by strain ATCC 25611 but not by clinical isolates lacking proliferative effects, and RT-qPCR confirmed higher *inpA* expression in OSCC-derived strains compared to non-cancer controls. Recombinant InpA (rInpA) mimicked the effect of CM, inducing proliferation and migration, which were blocked by PAR-2 inhibition. Transcriptomic and protein-level screening in SG cells revealed activation of EGFR and downstream RAS–RAF–MEK–ERK signaling. Furthermore, in a colorectal cancer (CRC) mouse model, oral administration of *P. intermedia* led to increased tumor formation, suggesting a broader oncogenic potential. These findings highlight InpA as a PAR-2–activating protease that may contribute to OSCC and potentially other cancers associated with oral microbiota dysbiosis.

## Introduction

Oral squamous cell carcinoma (OSCC) is a common malignancy worldwide and ranks as the fourth most prevalent cancer among men in Taiwan (1). It represents the most frequent form of head and neck cancer and can develop in various parts of the oral cavity, including the lips, tongue, cheeks, palate, and throat. The majority of oral cancers are squamous cell carcinomas, which are associated with poor prognosis and high mortality, with a 5-year survival rate of approximately 50%. Numerous factors contribute to OSCC development, such as gender, human papillomavirus (HPV) infection, alcohol consumption, tobacco use, betel nut chewing, poor oral hygiene, and alterations in the oral microbiota (2, 3). Notably, chronic periodontitis, a gum disease characterized by gingival inflammation and tooth loss, has been strongly linked to oral cancer (4). Poor oral hygiene facilitates the formation of biofilms, known as dental plaques, which contribute to oral microbiota dysbiosis. Anaerobic bacteria residing within these plaques produce acids and other virulence factors that contribute to periodontal inflammation and disease (5). Anaerobic bacteria residing within these plaques produce acids and other virulence factors that contribute to periodontal inflammation and disease.

Several bacterial genera, including *Porphyromonas, Fusobacterium*, and *Prevotella*, are found in higher abundance in patients with periodontitis compared to healthy individuals. (6, 7). *Porphyromonas gingivalis*, a Gram-negative anaerobe, is associated with periodontal disease and systemic conditions such as Alzheimer’s disease (8). Studies have also linked *P. gingivalis* to various cancers, including lung, pancreatic, prostate, colorectal cancer (CRC), and OSCC (9, 10). This bacterium produces multiple virulence factors, such as lipopolysaccharides (LPS), outer membrane vesicles (OMVs), capsules, and gingipains—cysteine proteases implicated in promoting tumor invasion and migration (11). Another oral bacterium, *Fusobacterium nucleatum*, another Gram-negative anaerobic bacterium, has been associated with CRC, breast, pancreatic, and oral cancers (12, 13). It secretes virulence factors including OMVs, LPS, and adhesins. The adhesin FadA interacts with E-cadherin to activate WNT/β-catenin signaling and promote cell proliferation (14). While Fap2 binds tumor-associated oligosaccharides and the immune checkpoint receptor TIGIT, potentially facilitating immune evasion (15). While much attention has focused on *P. gingivalis* and *F. nucleatum* in OSCC, the role of *Prevotella intermedia* remains poorly understood. *P. intermedia* is a Gram-negative, obligate anaerobic rod-shaped bacterium that requires vitamin K and hemin for growth and is associated with periodontal disease and acute necrotizing ulcerative gingivitis. Recent studies have reported elevated levels of *P. intermedia* in the oral microbiota of OSCC patients compared to healthy controls. Although this correlation suggests a potential role in oral carcinogenesis, the molecular contributions of this association remain unclear.

In this study, we demonstrate that *P. intermedia* strain ATCC 25611 promotes proliferation of normal oral epithelial cells (SG cell line) at a multiplicity of infection (MOI) of 100. Its conditioned medium (CM) also enhanced proliferation in SG, SCC-15, and SAS cell lines. Proteomic analysis using capillary electrophoresis/liquid chromatography-mass spectrometry (CE/LC-MS) revealed that a cysteine protease, interpain A (InpA), secreted by strain ATCC 25611, may be the critical factor driving this effect. In addition to stimulating proliferation, InpA also enhanced migration of OSCC cells. Furthermore, in a murine colorectal cancer model, colonization with *P. intermedia* increased tumor burden. These findings support a role for *P. intermedia*, and specifically InpA, in promoting OSCC development.

## Materials and methods

### Bacterial strains and cultures

*P. intermedia* strain ATCC 25611 was obtained from the Bioresource Collection and Research Center (BCRC, Taiwan), and culture conditions were adapted from a previously published protocol (16). Bacteria were grown anaerobically at 37 °C in supplemented Tryptic Soy Broth (ATCC 2722), consisting of tryptic soy broth (#211825, BD) enriched with 0.5% yeast extract (#212750, BD), 0.05% L-cysteine (#A3694, Applichem Panreac), 0.1% hemin (#H9039, Sigma), and 0.02% menadione (#M5625, Sigma). Clinical isolates of *P. intermedia* were obtained from the saliva of patients at National Cheng Kung University Hospital (NCKUH), under Institutional Review Board approval (IRB no. B-ER-106-234). Saliva samples were diluted in PBS and plated on Kanamycin-Vancomycin Laked Blood (KVLB) agar (TPM, Yilan, Taiwan). After 2–3 days of anaerobic incubation (5% CO_2_, 5% H_2_, 90% N_2_), black-pigmented colonies were selected. Colony identity was confirmed by Gram staining and PCR using *P. intermedia*-specific primers Pi-192 and Pi-486 (Supplementary Table 1). The PCR conditions were as follows: initial denaturation at 95 °C for 5 min; 35 cycles of 95 °C for 30 s, 55 °C for 30 s, and 72 °C for 30 s; and a final extension at 72 °C for 7 min.

### Cell lines and cultures

The immortalized human gingival keratinocyte, Smulow–Glickman (SG) cell (17), was kindly provided by Dr. Chuan-Fa Chang (NCKU) and cultured with Dulbecco’s Modified Eagle Medium (DMEM) (#11965-092, Gibco/Life Technology, Carisbad, CA, USA). Human tongue carcinoma cell lines, SAS cells (18), and SCC-15 cells (19) were obtained from Dr. Jhen-Wei Ruan (NCKU) and maintained in DMEM/F12 (D/F12) medium (#11330-032, Gibco). All media were supplemented with 10% FBS (#10437-028, Gibco) and 1% 100x Penicillin-Streptomycin-Amphotericin B solution (#CC501-0100, Simply Biotechnology, San Diego, CA, USA). Cells were incubated at 37 °C in a humidified atmosphere containing 5% CO_2_.

### Conditioned medium (CM) preparation

The preparation of *P. intermedia* condtioned medium was adapted from a previously published protocol (20). *P. intermedia* strains were anaerobically cultured in ATCC 2722 broth at 37 °C and were harvested when the OD600 value reached 1.0. The bacterial culture was centrifuged at 3220 xg for 3 min and discarded supernatant. The pellet was resuspended with a double volume of cell culture medium and anaerobically incubated at 37 °C for an additional 8 hr. Then the supernatant was collected after being centrifuged at 3220 xg for 10 min, filtered with a 0.22 µm filter (Merck Millipore, Burlington, MA, USA), and stored at 4 °C before use.

### Cell proliferation assay

The cell proliferation assay was performed as previously described (21). SG, SAS, or SCC-15 cells were dissociated by 0.05% trypsin-EDTA solution (#25300-054, Gibco) and resuspended in DMEM or D/F12 medium. A total of 1 × 10^4^ cells were seeded into each well of 24-well tissue culture plates (Corning, NY, USA) and treated with bacteria or CM treatment. CM was fractionated into four components using centrifugal filter devices with molecular weight cutoffs of 3 kDa, 10 kDa, and 30 kDa (Merck Millipore). Treatments included 10% CM; the PAR-2 antagonist FSLLRY-NH2 at 40 µM for SG cells or 20 µM for SCC-15 and SAS cells (#4751, TOCRIS/Bio-Techne, Minneapolis, MN, USA); and the cysteine protease inhibitor E64 at 5 µM for SG cells or 0.5 µM for SCC-15 and SAS cells (#324890, Merck Millipore). All treatments were administered in DMEM or D/F12 containing 10% FBS to a final volume of 1 ml. After incubation for 24, 48, or 72 hr at 37 °C in a 5% CO_2_ atmosphere, cells were washed once with 1× PBS and then dissociated using 0.05% trypsin-EDTA. The resulting cell suspension was mixed with an equal volume of 0.4% trypan blue (#15250-061, Gibco), and viable cells were counted using a Luna cell counter (Logos Biosystems, Gyeonggi-do, South Korea).

### Capillary electrophoresis/Liquid Chromatography-Mass spectrometry (CE/LC MASS) analysis

A total of 100 µg of protein in 60 µl rehydration buffer (7 M urea, 2 M thiourea, and 2% CHAPS) was mixed with 9.3 µl of 7.5% SDS (#75746, Sigma, Burlington, MA, USA) and 0.7 µl of 1 M dithiothreitol (#UR-DTT, UniRegion Bio-Tech, Taichung, Taiwan) prepared in 7 M urea and 0.01 M sodium acetate. The mixture was heated at 95 °C for 5 min, then placed on ice for 3 min. Next, 4 µl of 500 mM IAM (iodoacetamide in 50 mM ammonium bicarbonate) was added and incubated at room temperature for 30 min. Afterward, 52 µl of 50% trichloroacetic acid (TCA) (#100807, Merck Millipore) was added, and the solution was placed on ice for 15 min. The precipitate was collected by centrifugation at 15,871 ×g for 15 min at 4 °C, resuspended in 150 µl of 10% TCA, and centrifuged again at 15,871 ×g for 5 min at 4 °C. This washing step was repeated three times. The final pellet was resuspended in 100 µl of 100 mM ammonium bicarbonate and digested with 20 µl of trypsin solution (#V5111, Promega, Madison, WI, USA) at 37 °C for 16 hr. To inactivate trypsin, samples were stored overnight at −80 °C. Prior to analysis, samples were desalted using Ziptips (Merck Millipore). Protein identification was performed using CE/LC-MS on a Q Exactive Orbitrap Mass Spectrometer (Thermo Fisher Scientific).

### Interpain A gene expression quantitative analysis (RT-qPCR)

RNA extraction and RT-qPCR were adapted from previously described methods (22). A 2 ml culture of *P. intermedia* (OD_600_ = 0.8) was mixed with 4 ml of RNAprotect Bacteria Reagent (QIAGEN, Venlo, Netherlands). After centrifugation at 2,200 ×g for 5 min, the supernatant was discarded, and the pellet was stored at −80 °C until RNA extraction. RNA was purified using the RNeasy Mini Kit (QIAGEN), and residual DNA was removed using DNase (Promega). The resulting RNA was concentrated with the Clean & Concentrator Kit (ZYMO GENETICS, Seattle, WA, USA). cDNA was synthesized from the RNA using the SuperScript III First-Strand System with random hexamers (Invitrogen, Waltham, MA, USA). RT-qPCR was performed using GoTaq qPCR Master Mix (Promega) on a MyGo Pro real-time PCR instrument (IT-IS Life Science Ltd/DKSH, St Leonards, Australia) with primers inpA F, inpA R, Pi 16S rRNA F, and Pi 16S rRNA R (Supplementary Table 1). The thermal cycling conditions were as follows: initial denaturation at 95 °C for 10 min, followed by 40 cycles of denaturation at 95 °C for 15 s and annealing/extension at 60 °C for 1 min. A melt curve analysis was performed from 60 to 97 °C at a ramp rate of 0.1 °C/s. All procedures were conducted in accordance with the manufacturers’ instructions.

### InpA_N381_ expression vector construction and recombinant InpA_N381_ purification

Genomic DNA of *P. intermedia* ATCC 25611 was extracted using the MasterPure™ complete DNA and RNA purification kit (Lucigen Corp, Middleton, WI, USA) according to the manufacturer’s instructions. The gene region encoding the signal peptide, pro-domain, and catalytic domain (inpA_N381_) was amplified using a mixture of GoTaq G2 Flexi DNA Polymerase (Promega) and Pfu polymerase (Promega) with primers *inpA*_*N381*_ F and *inpA*_*N381*_ R (Supplementary Table1). The PCR product was digested with BamHI and XhoI and ligated into the pET21b(+) expression vector pre-digested with the same enzymes. The resulting plasmid, pET21b(+)::*inpA*_*N381*_, was verified by sequencing and chemically transformed into *Escherichia coli* C41 (DE3) cells. A single colony of *E. coli* C41 (DE3) harborin*g* pET21b(+)::*inpA*_*N381*_ was inoculated into 20 ml LB broth supplemented with ampicillin (100 µg/ml) and cultured at 37 °C for 14-16 hr with shaking at 170 rpm. The bacterial culture was then refreshed with 2 liters LB at the ratio of 1:100 and grew at 37 °C with shaking at 170 rpm until OD600 reached 0.4. Protein expression was induced by adding 0.2 mM isopropyl-β-D-thiogalactoside (IPTG; #101-367-93-1, Cyrusbioscience, Inc., Seattle, WA, USA), followed by incubation at 25 °C for an additional 16 hr with shaking (23). After induction, the bacterial culture was centrifuged at 3,300 ×g for 40 min at 4 °C. The resulting pellet was resuspended in 100 ml of 1× PBS and lysed by ultrasonication on ice (20 s pulse on, 100 s pulse off) until complete cell disruption. The lysate was then centrifuged at 16,000 ×g for 40 min at 4 °C. InpA_N381_ protein was purified from the supernatant using a HisTrap HP His-tag affinity column (Cytiva, Washington D.C., USA) according to the manufacturer’s instructions. The purified InpA_N381_ protein was washed with 100 volumes of 1× PBS and stored at −20 °C until use.

### Transwell assay

The Transwell assay was conducted as previously described (24). SAS or SCC-15 cells were resuspended in D/F12 medium supplemented with 10% FBS. Cell concentration was adjusted to 1 × 10^5^ cells/ml. A total of 100 µl of cell suspension was added to the upper chamber of Transwell inserts (8 µm pore size, Falcon/Corning, NY, USA) placed in a 24-well plate and incubated at 37 °C in 5% CO_2_. The lower chamber was filled with 10% CM, 20 µM FSLLRY-NH_2_, or 0.5 µM E64, in D/F12 containing 10% FBS, to a final volume of 600 µl. After 24 hr, cells in the upper chamber were removed with cotton swabs, and migrated cells on the lower side of the membrane were fixed with 70% ethanol for 10 min and stained with 0.2% crystal violet for 5 min. Five random fields per membrane were selected for cell counting under an inverted microscope (Leica Microsystems, Wetzlar, Germany).

### Wound healing assay

The wound healing assay was performed according to a previously reported method (24). SAS or SCC-15 cells were seeded in both compartments of each block insert (SPL Life Sciences Co., Gyeonggi-do, Korea), placed in 24-well plates, and incubated at 37 °C with 5% CO_2_ for 24 hr. The inserts were then removed, and the wells were washed once with 1× PBS. Cells were treated with 10% CM, 20 µM FSLLRY-NH_2_, or 0.5 µM E64, with the total volume adjusted to 1 ml using D/F12 containing 10% FBS. Images were captured every 2 hr using a microscope (OLYMPUS, Tokyo, Japan) to monitor gap closure. Gap widths were measured using ImageJ, and wound closure rates were calculated by dividing the change in width over time.

### Colorectal cancer (CRC) mouse model

The mouse model was modified from previously published protocols (25). Seven-week-old specific pathogen-free (SPF) male C57BL/6 mice were purchased from the Laboratory Animal Center of NCKU. Mice received a single intraperitoneal injection of azoxymethane (AOM; Sigma-Aldrich) at 12.5 mg/kg, followed by administration of 2% dextran sodium sulfate (DSS; MP Biomedicals, Irvine, CA, USA) in drinking water to induce colorectal cancer. Twenty mice were randomly assigned to four groups: MOCK, AOM/DSS, AOM/DSS + *F. nucleatum*, and AOM/DSS + *P. intermedia*. Mice underwent three cycles of DSS and bacterial treatment. Each cycle lasted 3 weeks: 5 days of 2% DSS in the first week, followed by two weeks of oral gavage with either 1× PBS or 1 × 10^9^ CFU of *F. nucleatum* (ATCC 23726) or *P. intermedia* (ATCC 25611) once per week. Mice were sacrificed at week 10, and colons were harvested, longitudinally opened, and tumor numbers were counted.

### Statistical analysis

Statistical analyses were conducted using GraphPad Prism 8 software. All data were obtained from experiments performed in triplicate. Comparisons between two groups were assessed using the Student’s t-test, while multiple group comparisons were analyzed by one-way ANOVA. Statistical significance was indicated as follows: *p* < 0.05 (*), *p* < 0.01 (**), *p* < 0.001 (***), and *p* < 0.0001 (****).

## Results

### *P. intermedia* ATCC 25611 enhances human oral cell proliferation

A previous study at National Cheng Kung University Hospital (NCKUH) reported a significant increase in *P. intermedia* abundance in patients with oral squamous cell carcinoma (OSCC) compared to healthy controls (6). To investigate the potential contribution of *P. intermedia* to OSCC development, we assessed its effect on oral epithelial cell proliferation. The reference strain ATCC 25611 was tested at multiple multiplicities of infection (MOIs) using the human normal oral epithelial cell line SG. After 24 hours of co-culture, *P. intermedia* ATCC 25611 significantly increased SG cell numbers at an MOI of 100 compared to the uninfected control group (Fig. 1A, 1B). In addition to ATCC 25611, we also examined six clinical isolates of *P. intermedia* obtained from patient saliva samples at NCKUH. Among these, strains H26219-3, H29383, and H29703 were isolated from OSCC patients, while strains H31082, H29823, and H31226 were obtained from individuals with non-cancerous oral conditions. SG cells were co-cultured with each clinical isolate at MOI 100. Unlike ATCC 25611, these strains did not induce a proliferation increase in SG cells; notably, strain H29383 even reduced SG cell numbers after 24 hours (Fig. 1B; Fig. S1A). We further found that only live *P. intermedia* was capable of promoting SG cell proliferation (Fig. 1C), likely due to its limited viability under aerobic conditions, which may confine its activity to the early co-culture period. Similarly, conditioned medium (CM) from strain ATCC 25611 increased SG cell proliferation, whereas CM from the clinical strains failed to elicit the same response (Fig. 1D; Fig. S1B). These findings were corroborated using EdU incorporation assays (Fig. S1C), which supports the role of a secreted factor in enhancing proliferation. To identify this active factor, we subjected the CM to heat inactivation, which abolished its proliferative effect (Fig. 1D), indicating the involvement of a heat-sensitive, likely proteinaceous molecule. Further size-fractionation of CM revealed that only the >30 kDa fraction retained stimulatory activity (Fig. 1E). Additionally, CM from ATCC 25611 significantly enhanced proliferation in OSCC cell lines SCC-15 and SAS after 48 and 72 hours, respectively (Fig. 1F, 1G), suggesting that *P. intermedia* promotes oral epithelial cell proliferation through the action of one or more secreted proteins.

**Figure 1.**
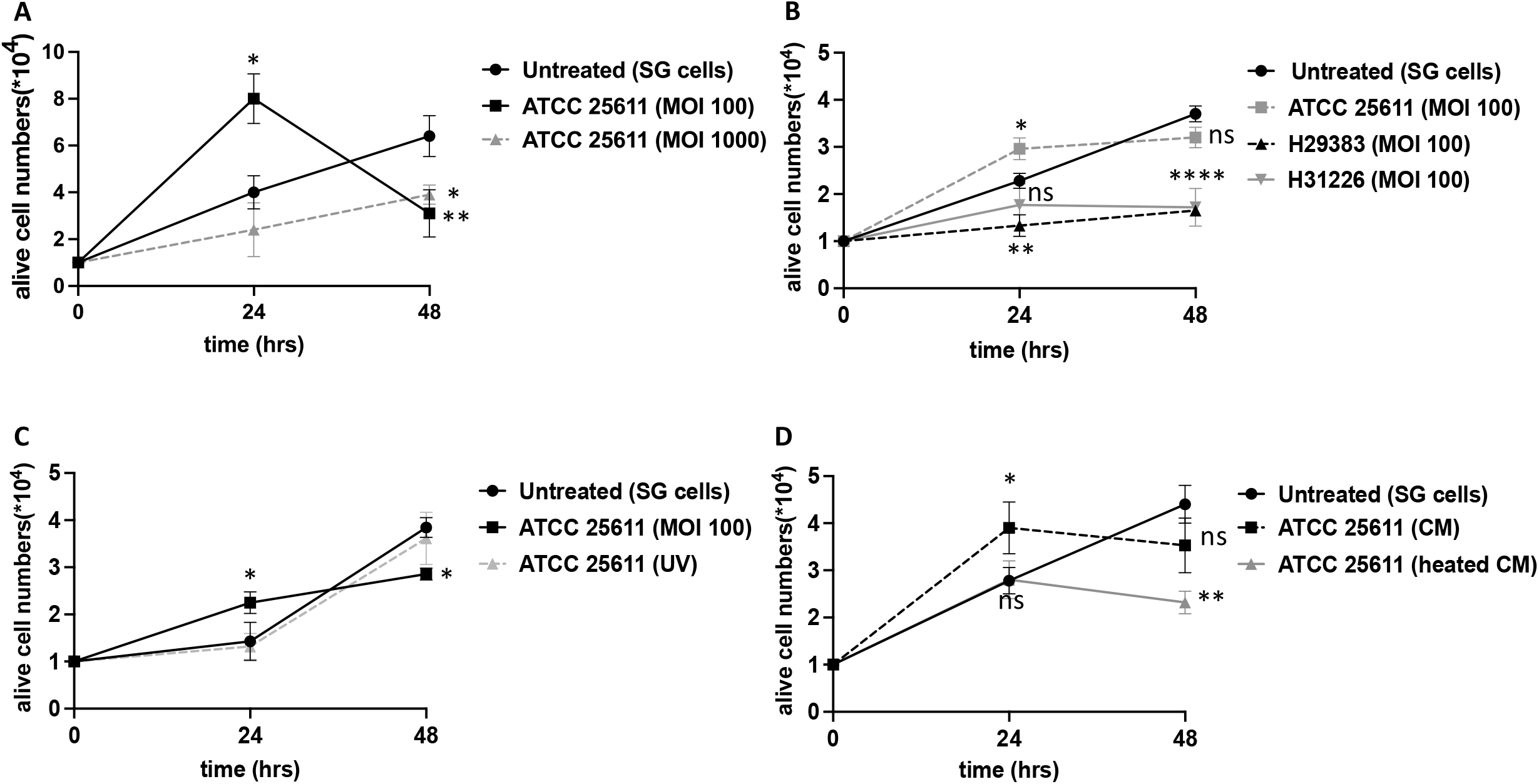
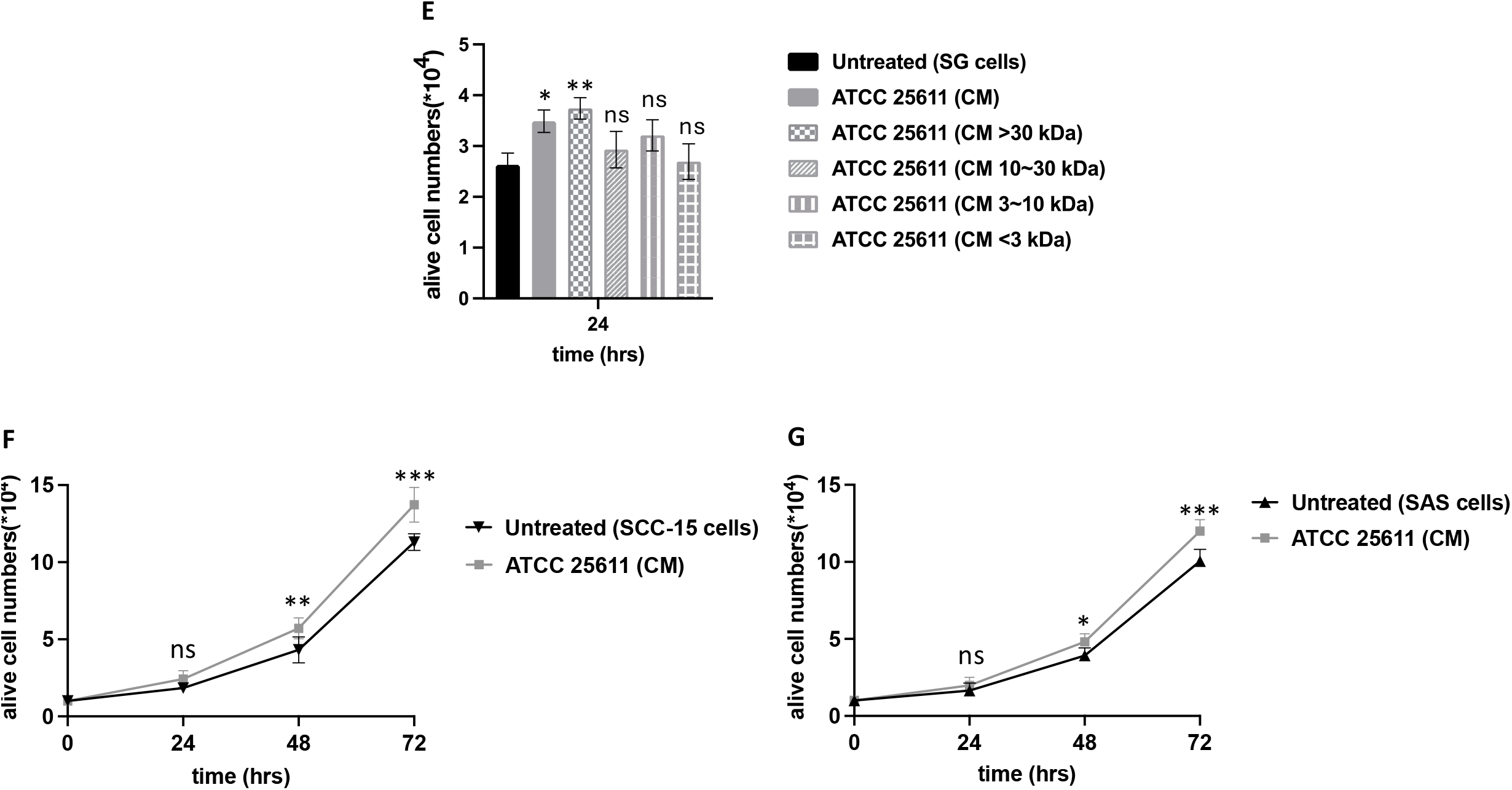
*P. intermedia* ATCC 25611 enhances proliferation of normal and cancerous oral epithelial cells. (A) Human normal oral epithelial SG cells were infected with live *P. intermedia* strain ATCC 25611 at a multiplicity of infection (MOI) of 100 or 1000 for 24 or 48 hours. (B) SG cells were co-cultured with strain ATCC 25611 or clinical isolates H29383 and H31226 at MOI 100. (C) Comparison of live versus UV-inactivated *P. intermedia* strain ATCC 25611 on SG cell proliferation. (D) Proliferative effect of conditioned medium (CM) from ATCC 25611 versus heat-inactivated CM on SG cells. (E) CM of strain ATCC 25611 was fractionated by molecular weight using centrifugal filters: >30 kDa, 10–30 kDa, 3–10 kDa, and <3 kDa. (F–G) CM from ATCC 25611 was applied to oral squamous carcinoma cell lines SCC-15 and SAS for 24, 48, and 72 hours. All results are representative of three independent experiments. Data are shown as mean ± SD. Statistical analysis was performed by one-way ANOVA (A–C) or t-test (D). ns: not significant; \**p* < 0.05; \*\**p* < 0.01; \*\*\**p* < 0.001; \*\*\*\**p* < 0.0001.

### Interpain A plays a centrol role in *P. intermedia* ATCC 25611 promoting oral cell proliferation

Following our observation that strain ATCC 25611 and several clinical isolates exhibited divergent effects on SG cell proliferation, we hypothesized that these differences could be attributed to the expression of specific secreted protein(s). To investigate this, we conducted proteomic analysis of conditioned medium (CM) from strain ATCC 25611 and the non-proliferative strain H31226 using capillary electrophoresis/liquid chromatography-mass spectrometry (CE/LC-MS). Several proteins were found to be more abundant in the CM of ATCC 25611 compared to that of H31226 (Supplementary Table 2). Among these, interpain A (InpA), a secreted cysteine protease, was particularly enriched—present at levels thirteen times higher in ATCC 25611 than in H31226 (Fig. 2A). InpA is a 90 kDa cysteine protease previously reported to facilitate heme acquisition for bacterial growth and to degrade complement factor C3, aiding immune evasion (26-28). Notably, *P. gingivalis*, another oral anaerobe strongly associated with chronic periodontitis and cancer, produces similar cysteine proteases called gingipains, which have been shown to activate protease-activated receptors (PARs) on oral epithelial cells (11). Gingipains, in particular, have been implicated in OSCC progression through activation of PAR-2 signaling, enhancing cell migration and invasiveness (29, 30). Given these functional similarities, we hypothesized that InpA may enhance oral epithelial cell proliferation via activation of the PAR-2 signaling pathway. To test this hypothesis, SG cells were treated with CM from ATCC 25611 in the presence of either E64, a cysteine protease inhibitor, or FSLLRY-NH_2_, a PAR-2 antagonist. After 24 hours, the proliferation-promoting effects of CM were significantly attenuated by both inhibitors (Fig. 2B). A similar inhibition pattern was observed under direct bacterial co-culture conditions. These results were consistent across SCC-15 and SAS oral cancer cell lines, where the proliferative effects observed after 48 hours were also blocked by E64 or FSLLRY-NH_2_ treatment (Fig. 2C, 2D). To further confirm InpA’s direct effect, we expressed and purified the recombinant catalytic domain of InpA (InpA_N381_) in *E. coli* strain C41. InpA_N381_ stimulated SCC-15 proliferation in a manner consistent with that of native CM, and again, this effect was neutralized by either E64 or FSLLRY-NH_2_ treatment (Fig. 2E). Importantly, treatment with E64 or FSLLRY-NH_2_ alone had no effect on the proliferation of SG, SCC-15, or SAS cells (Fig. S2), confirming that the inhibitors’ effects were specific to the activity of InpA or the associated pathway. We next investigated whether differences in InpA production could explain the observed strain-dependent variability in proliferation. RT-qPCR analysis revealed significantly higher *inpA* expression in ATCC 25611 compared to all clinical isolates tested (Fig. 2F). Furthermore, among the clinical strains, those isolated from OSCC patients (H26219-3, H29383, and H29703) exhibited higher *inpA* expression than strains from patients with non-cancerous oral diseases (H31082, H29823, and H31226). These findings collectively indicate that *P. intermedia* promotes oral epithelial cell proliferation via InpA and the PAR-2 signaling axis and that this activity is strain-dependent, correlating with InpA production levels.

**Figure 2.**
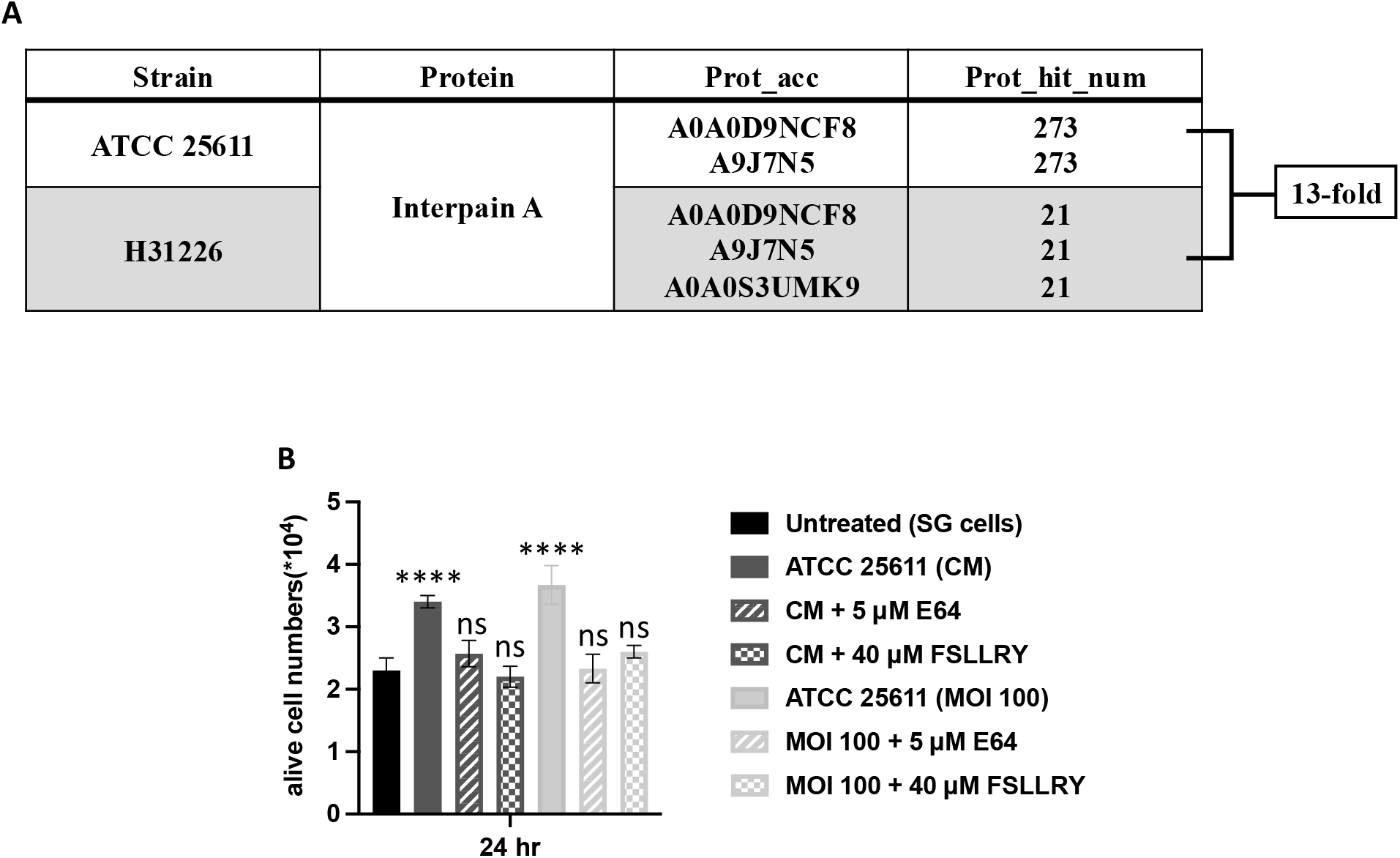
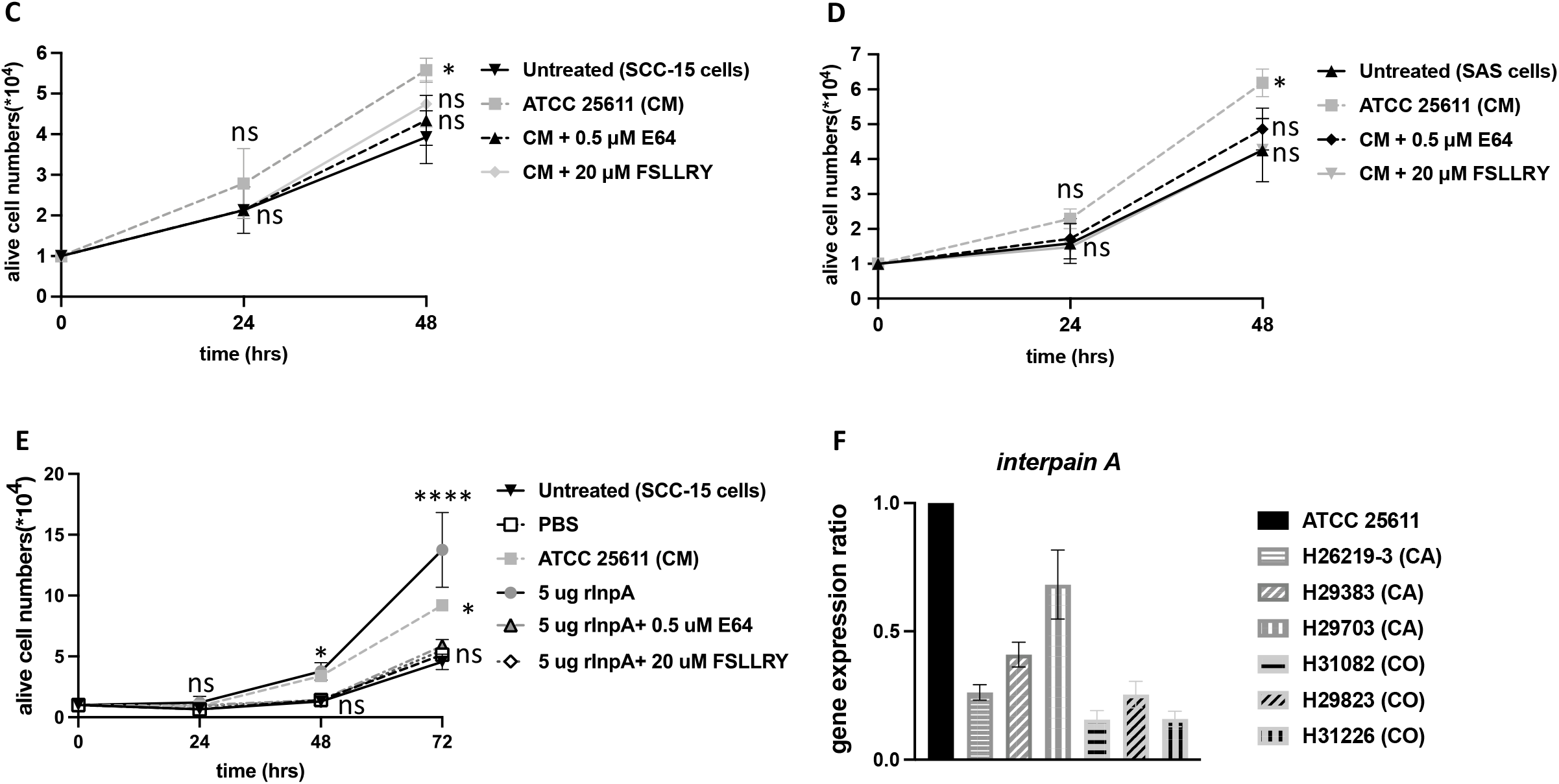
InpA promotes oral cell proliferation via PAR-2 activation. (A) CE/LC-MS identified elevated levels of InpA in strain ATCC 25611 compared to clinical strain H31226, which did not stimulate SG cell proliferation. (B) SG cell proliferation in response to CM or live *P. intermedia* (MOI 100), with or without 5 µM E64 (protease inhibitor) or 40 µM FSLLRY-NH_2_ (PAR-2 antagonist). (C-D) SCC-15 and SAS cell proliferation following treatment with CM from ATCC 25611, with or without 0.5 µM E64 or 20 µM FSLLRY-NH_2_. (E) Five micrograms of recombinant InpA (rInpA) were applied to SCC-15 cells, with or without E64 or FSLLRY-NH_2_. PBS served as a vehicle control; CM from ATCC 25611 served as a positive control. (F) Quantitative RT-PCR analysis of *inpA* gene expression in ATCC 25611 and six clinical isolates. Strains H26219-3, H29383, and H29703 were from oral cancer patients (CA), while H31082, H29823, and H31226 were from control patients (CO). Expression levels were calculated using the 2^–ΔΔCt method. All results are representative of three independent experiments. Data are expressed as mean ± SD and analyzed by one-way ANOVA. ns: not significant; \**p* < 0.05; \*\*\*\**p* < 0.0001.

### InpA-Mediated Promotion of OSCC Cell Migration

During cancer progression, tumor cells initially undergo uncontrolled growth at the primary site, followed by invasion and dissemination to distant organs—a process known as metastasis. Metastasis involves the degradation of the extracellular matrix, intravasation into the vasculature, and colonization of secondary tissues, and is a major contributor to cancer-related mortality (31). Given our findings that *P. intermedia* promotes oral epithelial cell proliferation, we next investigated whether it also influences OSCC cell migration. To address this, we employed two complementary assays—wound healing and transwell migration assays—using SCC-15 and SAS OSCC cell lines. In the wound healing assay, treatment with conditioned medium (CM) from *P. intermedia* strain ATCC 25611 significantly enhanced the migration rate of both cell lines at 2 and 4 hours post-treatment compared to untreated controls (Fig. 3A, 3B). To determine whether this migration-promoting effect involved InpA and PAR-2 signaling, we co-treated cells with either E64 or FSLLRY-NH_2_. Both inhibitors effectively abrogated the CM-induced enhancement of cell migration. Similarly, in the transwell migration assay, CM from ATCC 25611 significantly increased the number of migrating SCC-15 and SAS cells after 24 hours (Fig. 3C, 3D). Treatment with E64 or FSLLRY-NH_2_ had no significant effect on SCC-15 and SAS cell migration in either wound healing or transwell assays (Fig. S3). Co-treatment with E64 or FSLLRY-NH_2_ suppressed this enhanced migratory response. Furthermore, recombinant InpA_N381_ purified from *E. coli* elevated SCC-15 cell migration (Fig. 3E), and this effect was also blocked by both E64 and FSLLRY-NH_2_. Together, these findings suggest that *P. intermedia* facilitates OSCC cell migration through an InpA-dependent mechanism involving PAR-2 signaling, paralleling its role in cell proliferation.

**Figure 3.**
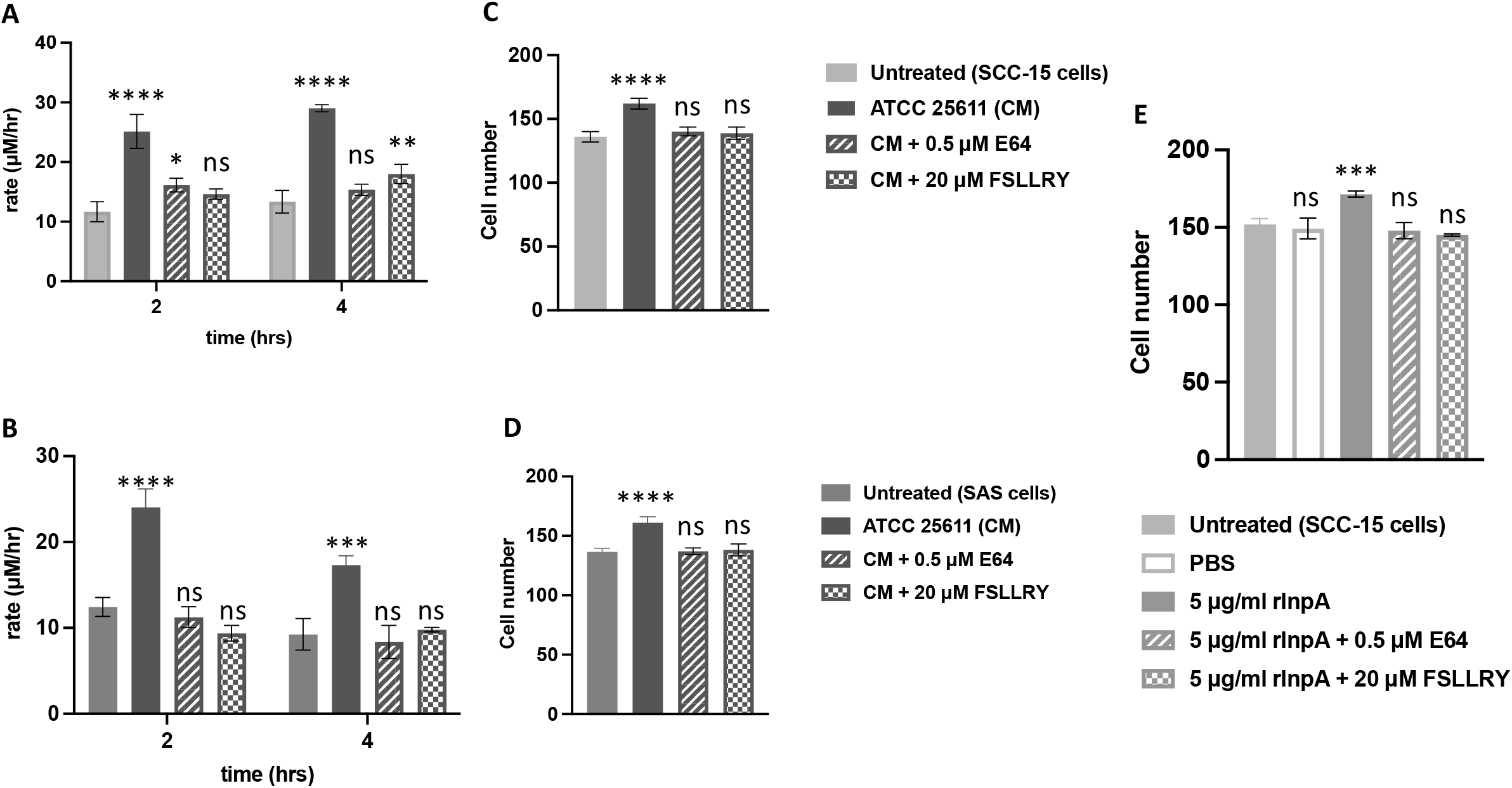
InpA enhances OSCC cell migration through PAR-2 signaling. (A–B) Wound healing assays in SCC-15 (A) and SAS (B) cells treated with standard medium (D/F12), CM from ATCC 25611, or CM with 0.5 µM E64 or 20 µM FSLLRY-NH_2_. Cell migration was assessed at 2 and 4 hours and calculated as migration rate (µm/hr). (C–D) Transwell migration assays in SCC-15 (C) and SAS (D) cells treated as above for 24 hours. Migrated cells were quantified under a phase-contrast microscope. (E) Transwell migration of SCC-15 cells treated with 5 µg/mL recombinant InpA (rInpA) with or without E64 or FSLLRY-NH_2_. All data represent triplicate experiments and are shown as mean ± SD. One-way ANOVA was used for statistical analysis. ns: not significant; \*\**p* < 0.01; \*\*\**p* < 0.001; \*\*\*\**p* < 0.0001.

### *P. intermedia* increases tumor burden in a colorectal cancer mouse model

An increasing body of evidence suggests that oral microbiota not only contributes to oral diseases but also plays a role in systemic conditions such as cystic fibrosis, cardiovascular disease, rheumatoid arthritis, and gastrointestinal disorders (32). Furthermore, several oral bacteria have been linked to cancers in distal organs, including the pancreas, breast, stomach, esophagus, and colon (32). The concept of the oral-gut axis proposes that oral bacteria can translocate to the gastrointestinal tract, where they may disrupt the gut microbial balance and contribute to diseases such as inflammatory bowel disease and colorectal cancer (CRC) (33, 34). Notably, *Fusobacterium nucleatum*, an oral anaerobe, has been identified as an important contributor to CRC progression (35). Given the pro-tumorigenic effects of *P. intermedia* observed in OSCC, we hypothesized that it may also influence CRC development. To test this, we used an azoxymethane (AOM) and 2% dextran sulfate sodium (DSS)-induced CRC model in C57BL/6 mice, followed by oral gavage with either *F. nucleatum* or *P. intermedia*. Compared to the AOM/DSS-only group, both bacterial treatments significantly increased tumor formation and elevated the number of colonic tumors (Fig. 4B, 4C). These results suggest that *P. intermedia*, like *F. nucleatum*, may contribute to CRC pathogenesis and should be considered a relevant microbial factor in colorectal tumorigenesis.

**Figure 4.**
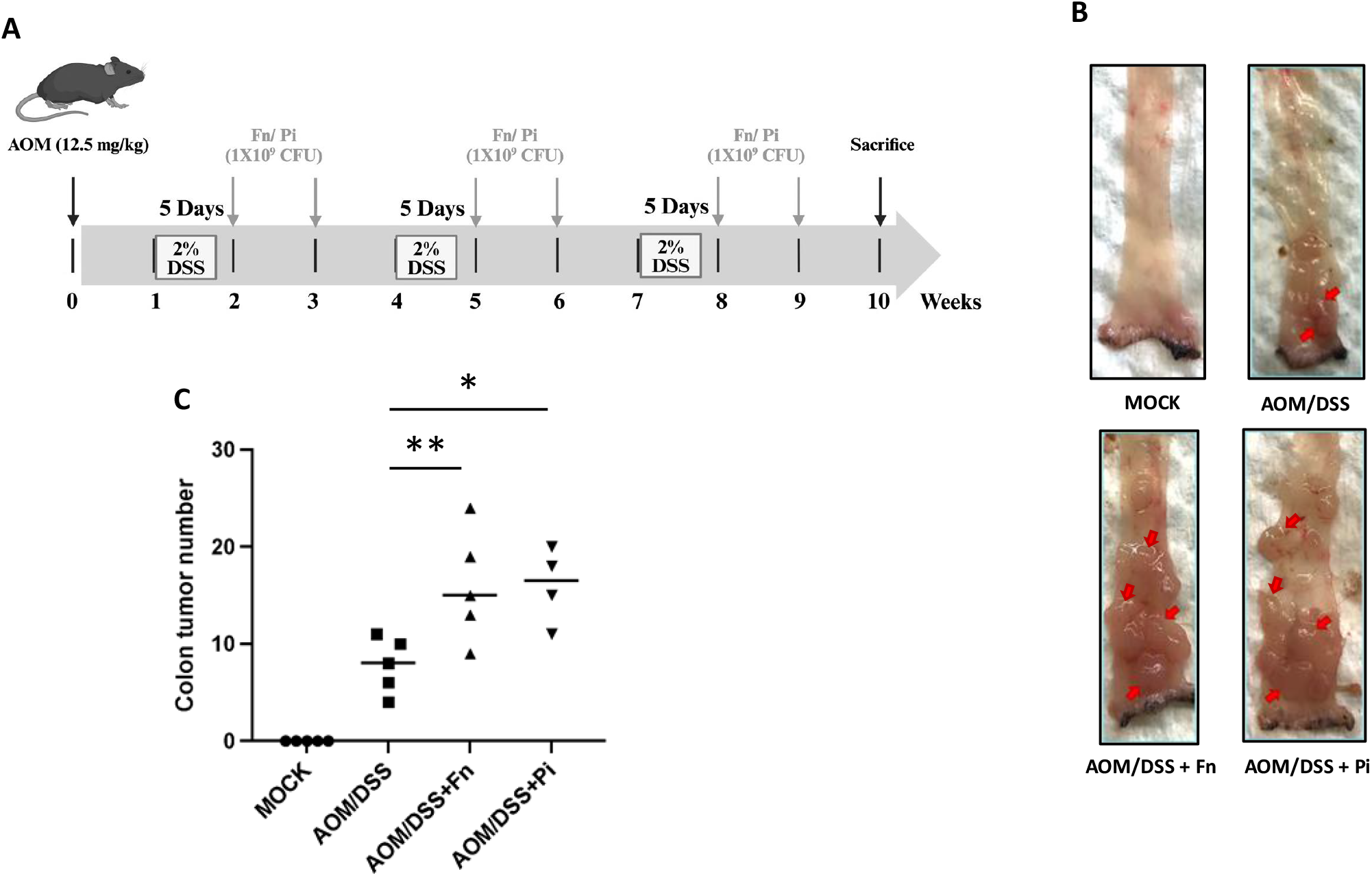
Oral administration of *P. intermedia* increases colorectal tumor burden in mice. (A) Schematic diagram of the azoxymethane (AOM)/dextran sulfate sodium (DSS)-induced colorectal cancer (CRC) mouse model. Mice received a single intraperitoneal injection of AOM (12.5 mg/kg), followed by three cycles of 2% DSS in drinking water (5 days per cycle, separated by 16 days of regular water). During the DSS cycles, 1 × 10^9^ CFU of *P. intermedia* ATCC 25611 or *F. nucleatum* ATCC 23726 was administered via oral gavage. Mice were euthanized at week 10. (B) Representative images of distal colon morphology from AOM/DSS mice with or without bacterial infection. (C) Quantification of tumor number in the distal colon. Each group included five mice. ns: not significant; \**p* < 0.05.

## Discussion

The present study demonstrated that *P. intermedia* may play a significant role in oral cancer development. Interpain A (InpA), a cysteine protease secreted by *P. intermedia*, functioned as an effector that enhanced both oral epithelial cell proliferation (Fig. 2B) and OSCC cell migration (Fig. 3A–D). Importantly, these effects were linked to the protease-activated receptor-2 (PAR-2) signaling pathway activation. Prior studies examining salivary samples from OSCC patients reported increased expression of various oral bacterial virulence genes, including *fadA* and *inpA* (36). Notably, PAR-2 has been implicated in promoting cell proliferation, metastasis, and invasion in multiple cancers, including OSCC, esophageal carcinoma, and pancreatic cancer (37). Previous studies have also shown that gingipains, secreted cysteine proteases from *P. gingivalis*, can activate the PAR-2/NF-κB signaling axis and induce proMMP9 expression, thereby facilitating OSCC invasion (11). These findings are consistent with our observations implicating InpA and PAR-2 signaling in OSCC progression.

Our results showed that *P. intermedia*-CM treatment promoted proliferation in both normal oral keratinocytes (SG cells) and OSCC cell lines (SCC-15 and SAS) (Fig. 1D–G). Interestingly, the proliferative effect manifested at different time points between cell types: SG cells responded within 24 hours, whereas SCC-15 and SAS cells showed continued stimulation at 48 and 72 hours. Moreover, SG cell numbers decreased after 48 hours, suggesting that normal oral cells may retain mechanisms to counterbalance proliferative signals, while cancer cells have lost this regulatory capacity. A previous study of *P. gingivalis* reported G_1_ cell cycle arrest and apoptosis in trophoblasts, as well as in osteoblastic/stromal cells following gingipain exposure (38, 39). Gingipains have also been shown to induce apoptosis in oral epithelial cells (40). It is possible that InpA exerts similar effects in SG cells following initial proliferation, potentially through the regulation of cell cycle or apoptotic pathways. This observation warrants further investigation. Our findings also highlight the need to clarify whether InpA differentially activates downstream PAR-2 signaling in normal versus malignant oral cells.

The concept of an oral-gut microbiome axis is increasingly supported by evidence showing that oral dysbiosis can influence both local and systemic health (41); A growing number of studies have linked oral microbiota imbalances with gastrointestinal diseases, including colorectal cancer (CRC) (42). Recent findings suggest that *P. intermedia* enhances CRC cell proliferation and migration through secreted protein effectors, similar to those identified in our OSCC model (43). The promoting effect was also from *P. intermedia-*secreted protein(s). Those results were similar to our findings (Fig. 2C-E). In agreement with these findings, our data showed that oral administration of *P. intermedia* increased tumor burden in an AOM/DSS-induced CRC mouse model (Fig. 4). These results reinforce the idea that *P. intermedia* may act as a tumor-promoting microbe across multiple tissue contexts. Beyond cancer, *P. intermedia* and other oral pathogens have been implicated in the pathogenesis of various systemic diseases (44). Oral bacteria are capable of releasing inflammatory mediators that may affect the central nervous system. Elevated serum antibody levels against *P. gingivalis, F. nucleatum*, and *P. intermedia* have been observed in patients with Alzheimer’s disease (45). Gingipains from *P. gingivalis* have been detected in the brains of Alzheimer’s patients and are known to increase blood–brain barrier permeability (8, 46). Moreover, gingipains have been associated with other neurodegenerative diseases, such as Parkinson’s disease (47). Periodontal bacteria have also been identified in atheromatous plaques and are believed to contribute to cardiovascular disease pathogenesis (48, 49). These findings position *P. intermedia* as a potentially important contributor to systemic inflammatory and degenerative diseases. Our study also supports the notion that the pro-proliferative effects of *P. intermedia* are strain-dependent. In SG cells, only strain ATCC 25611 induced proliferation at MOI 100 (Fig. 1B, S1A), and only its conditioned medium promoted proliferation (Fig. 1D, S1B). Proteomic analysis confirmed that this strain secreted significantly higher levels of InpA than clinical isolates such as H31226 (Fig. 2A), and RT-qPCR revealed higher *inpA* expression in strains isolated from OSCC patients compared to non-cancer controls (Fig. 2F). These observations raise the possibility of using salivary InpA levels as a biomarker for OSCC risk or diagnosis. Although there was no statistically significant difference between clinical strain H26219-3 or H29383 and the untreated control group at the 24-hour time point (Fig. S1B), SG cell proliferation showed a mild upward trend in response to both strains isolated from OSCC patients. This observation suggests that these strains may exert weak but measurable biological effects. Given that such bacteria can persistently colonize the oral cavity, it is plausible that their cumulative impact over time could become biologically relevant. However, short-term cell-based assays are unlikely to capture such long-term effects.

To explore the mechanism underlying InpA-dependent proliferation, we performed a Micro-Western Array focusing on signaling pathways related to cell growth in SG cells (Fig. S4). We found that the EGFR signaling pathway was upregulated, with RAF expression increasing at 8 hours and decreasing at 24 hours. This aligns with prior studies showing that the EGFR–RAS–RAF–MEK–ERK axis is frequently upregulated in OSCC (50). While these findings provide a preliminary mechanistic link between InpA and cancer signaling pathways, further validation is required.

In conclusion, our study demonstrates that InpA, a cysteine protease secreted by *P. intermedia*, plays a pivotal role in promoting oral epithelial cell proliferation and OSCC cell migration. These effects appear to be mediated via the PAR-2 signaling pathway and are strain-dependent. The results suggest a potential microbial mechanism contributing to OSCC development and lay the groundwork for future investigations into therapeutic strategies targeting microbial virulence factors.

## Acknowledgements

We thank the Laboratory Animal Center, College of Medicine at National Cheng Kung University (NCKU), Taiwan, accredited by AAALAC International and Taiwan Animal Consortium, for their support in caring for mice. We appreciate the participation of all patients in this study. We are grateful to all the unmentioned clinicians and recruiters. We also thank Dr. Chuan-Fa Chang and Dr. Jhen-Wei Ruan provided the cell lines.

## Financial support

This work was supported by the National Science and Technology Council (Grants: 113-2314-B-006-043-MY3) awarded to J. W. C. and OSU-CHS Startup awarded to I.H.H.. The funding agencies played no role in study design, data collection or analysis, manuscript preparation, or decision to publish.

## Authors’ contributions

S.-M. C., Y.-J. C., W.-R. W. and N.-H. W. performed the experiments. S.-M. C. wrote this manuscript. I-H. H. assisted with the experimental design and reviewed this manuscript. C.-P. C. performed the microwestern array experiments and reviewed this manuscript. J.-R. H., J. S C., and J.-Y. C. provided clinical patients’ saliva. J.-W. C. contributed to design, provide the concept, and supervise this manuscript. All authors have read and approved the final version of this manuscript.

## Notes

**Competing Interest Statement:** The authors declare no potential conflicts of interest.

### Competing Interest Statement

The authors have declared no competing interest.

### Summary of Updates

The content has been revised. Figures have also been revised.

