## Supplementary material for "The protease interpain A of *Prevotella intermedia* promotes human oral squamous cell carcinoma cells proliferation and migration": Fig S

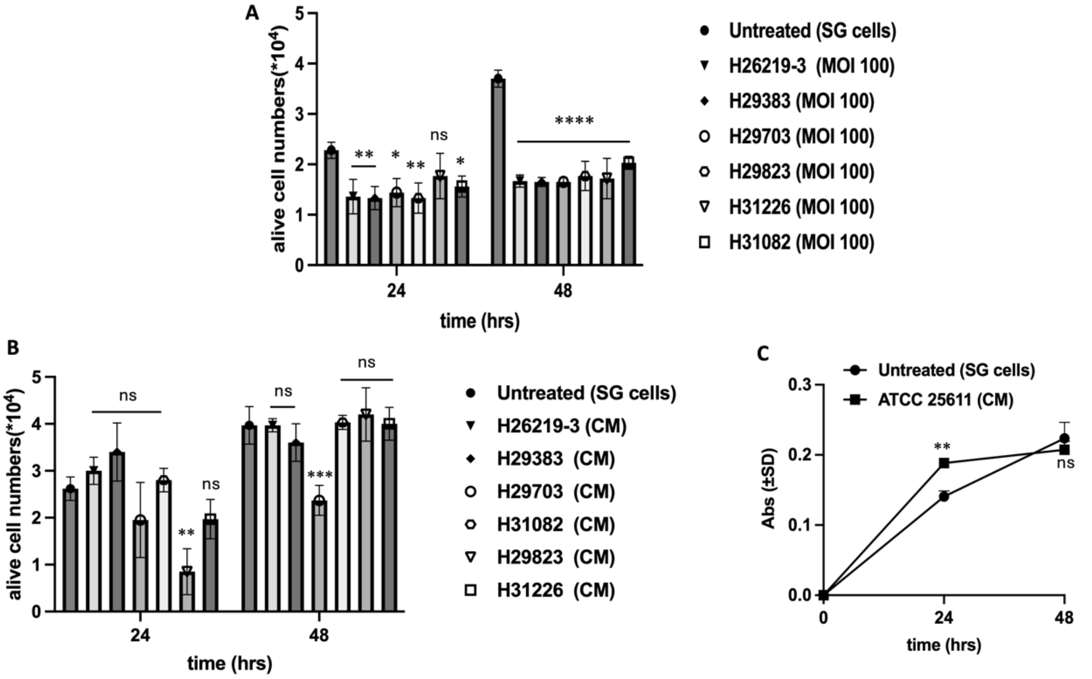


**Figure S1. Clinical *P. intermedia* strains do not significantly increase SG cell proliferation.** (A) Proliferation of SG cells treated with six clinical *P. intermedia* strains at MOI 100. Case strains: H26219-3, H29383, and H29703. Control strains: H29823, H31226, and H31082. (B) Proliferation of SG cells treated with 10% conditioned medium (CM) from the six clinical *P. intermedia* strains. (C) EdU proliferation assay of SG cells treated with CM from *P. intermedia* strain ATCC 25611. All results represent three independent experiments. Data are presented as mean ± SD (A & B) or absorbance ± SD (C). Statistical analysis was performed using one-way ANOVA (A & B) or unpaired t-test (C). ns: not significant; *$p < 0.05$; **$p < 0.01$; ***$p < 0.001$; ****$p < 0.0001$.

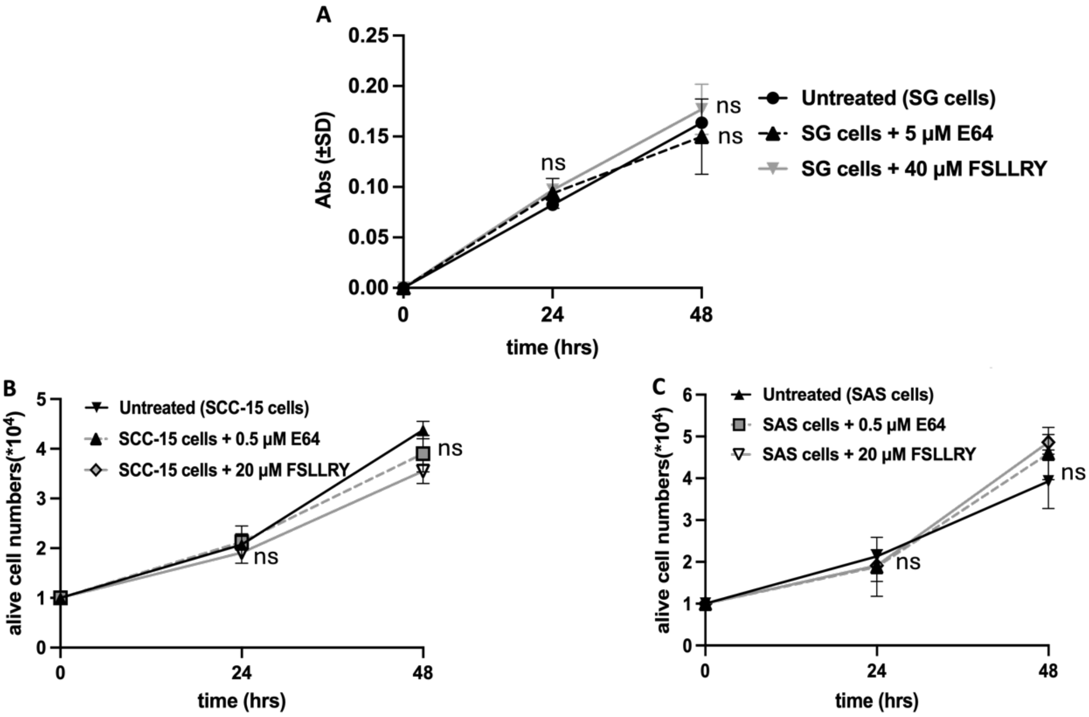


**Figure S2. Neither cysteine protease inhibitor (E64) nor PAR-2 receptor antagonist (FSLLRY-$NH_2$) affects the proliferation of SG, SCC-15, or SAS cells.** (A) EdU proliferation assay of SG cells treated with 5 μM E64, or 40 μM FSLLRY-$NH_2$. Proliferation of SCC-15 (B) and SAS (C) cells treated with or without 0.5 μM E64 or 20 μM FSLLRY-$NH_2$. All results represent three independent experiments. Data are presented as absorbance ± SD (A) or mean ± SD (B, C). Statistical analysis was performed using one-way ANOVA. ns: not significant; *$p < 0.05$.

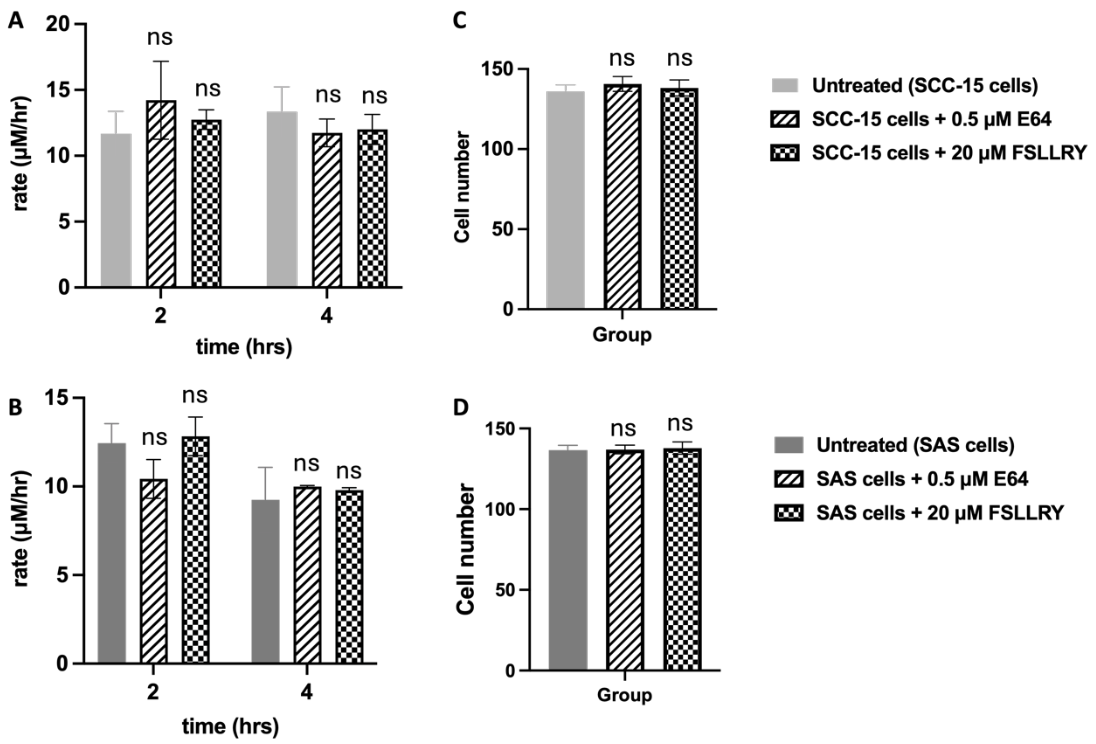


**Figure S3. E64 and FSLLRY-$NH_2$ alone do not affect the migration of SCC-15 or SAS cells.** Wound healing assay of SCC-15 (A) and SAS (B) cells treated with 0.5 μM E64 or 20 μM FSLLRY-$NH_2$. Transwell migration assay of SCC-15 (C) and SAS (D) cells treated under the same conditions. Data are presented as mean ± SD. All results represent triplicate experiments. Statistical analysis was performed using one-way ANOVA. ns: not significant.

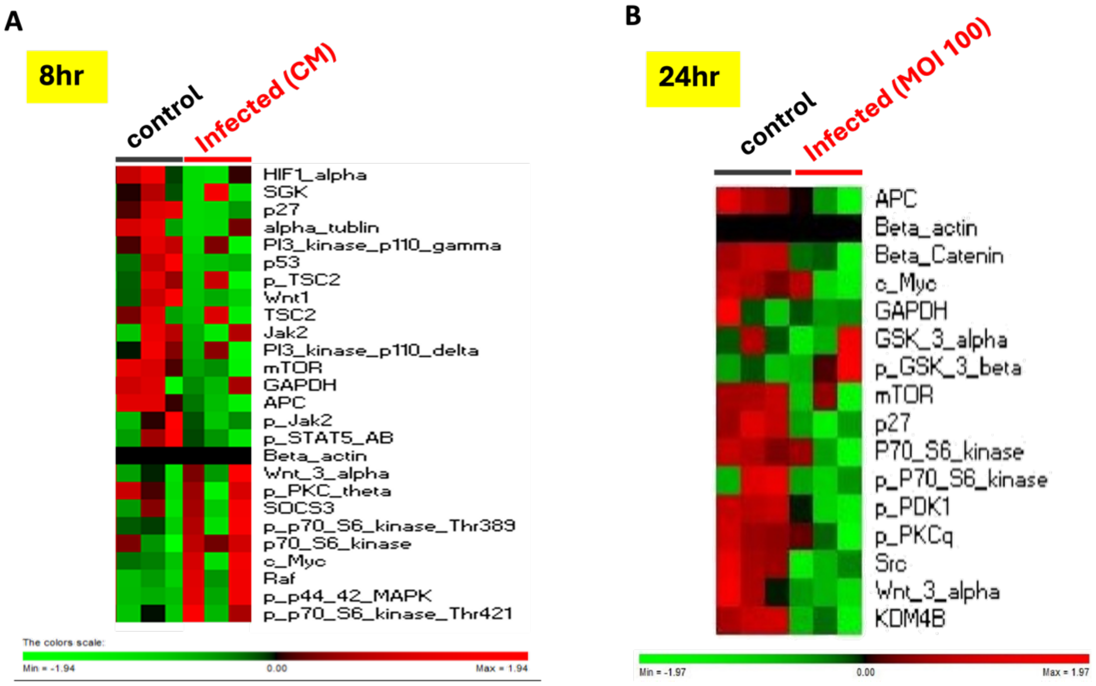


**Figure S4. Identification of cell proliferation pathways induced by *P. intermedia* in OSCC progression using Microwestern Array.** SG cells (1 × $10^5$ per well) were seeded in 24-well plates and incubated with 10% CM from *P. intermedia* strain ATCC 25611 for 8 hr or with bacteria at MOI 100 for 24 hr. After washing three times with PBS, cells were dissociated with 0.05% trypsin-EDTA, centrifuged at 395 × g, and washed twice with PBS. Samples were loaded onto SDS-PAGE arrays using a non-contact microarray printer. Each 96-well array contained three control samples, three infected samples (10% CM or MOI 100), and one marker. Heatmaps were generated using PermutMatrixEN software, showing proteins with expression levels >1.5-fold relative to β-actin. (A) Microwestern analysis of SG cells treated with 10% CM for 8 hr.(B) Microwestern analysis of SG cells infected at MOI 100 for 24 hr. Increased expression is shown in red; decreased expression in green.

**Table 1. Bacterial strains, cell lines and primers used in this study.**

| | Description | Source or Reference |
|---|---|---|
| **Bacterial strain** | | |
| ATCC 25611 | *P. intermedia* strain | BCRC† |
| H26219-3 | *P. intermedia* clinical isolated strain | NCKUH* |
| H29383 | *P. intermedia* clinical isolated strain | NCKUH* |
| H29703 | *P. intermedia* clinical isolated strain | NCKUH* |
| H31082 | *P. intermedia* clinical isolated strain | NCKUH* |
| H29823 | *P. intermedia* clinical isolated strain | NCKUH* |
| H31226 | *P. intermedia* clinical isolated strain | NCKUH* |
| ATCC 23726 | *F. nucleatum* strain | BCRC† |
| *E. coli* C41(DE3)-pET21b (+)::*inpA*$_{N381}$ | *E. coli* C41(CE3) strain carrying pET21b plasmid with *inpA*$_{N381}$ | This study |
| **Cell lines** | | |
| Smulow–Glickman (SG) | human gingival epithelial cell | Dr. Chuan-Fa Chang (NCKU) |
| SCC-15 | human tongue squamous cell carcinoma | Dr. Jhen-Wei Ruan (NCKU) |
| SAS | human tongue squamous cell carcinoma | Dr. Jhen-Wei Ruan (NCKU) |
| **Primers (5' to 3')** | | |
| **PCR** | | |
| Pi-192 | CCACATATGGCATCTGACGTGGAC | Reference |
| Pi-486 | CCCGCTTTACTCCCCAACAA | Reference |
| *inpA* F | TTTCAAGTATCCAGTGCGAGG | This study |
| *inpA* R | ATATTTGCCCAGTCGTAGGTG | This study |
| **qRT-PCR** | | |
| Pi 16s rRNA F | ATATGGCATCTGACGTGGAC | This study |
| Pi 16s rRNA R | ACAAGCTAATCAGACGCATCCCCATC | This study |
| **pET21b (+)::*inpA*$_{N381}$ construction** | | |
| *inpA*$_{N381}$ F | C**GGATCC**ATGAAAATTAAACAAAACCTATTCGTAGCCTTTGTG | This study |
| *inpA*$_{N381}$ R | C**CTCGAG**TGGTTTTCCGTAAACACCTCTAACCATATC | This study |

†BCRC: Bioresource Collection and Research Center

*NCKUH: National Cheng Kung University Hospital

**Supplementary Table 1: Bacterial strains, cell lines and primers used in this study.**

**Table 2: Proteins with 2-fold higher in strain ATCC 25611 than in strain H31226 by CE/LC MASS.**

| Protein | ATCC 25611-prot_hit_num | H31226-prot_hit_num | Fold |
|---|---|---|---|
| Succinate dehydrogenase | 127 | 6 | 21.2 |
| Membrane protein | 262 | 13 | 20.2 |
| NigD-like protein | 419 | 31 | 13.5 |
| **Interpain A** | **273** | **21** | **13.0** |
| DNA-binding protein | 501 | 57 | 8.8 |
| 50S ribosomal protein L7/L12 | 85 | 10 | 8.5 |
| Fumarate reductase/succinate dehydrogenase flavoprotein subunit | 46 | 6 | 7.7 |
| Succinate dehydrogenase flavoprotein subunit | 46 | 6 | 7.7 |
| Aspartate--tRNA ligase | 164 | 41 | 4.0 |
| ABC transporter ATP-binding protein | 411 | 111 | 3.7 |
| TPR domain protein | 293 | 82 | 3.6 |
| Transcriptional regulator | 388 | 112 | 3.5 |
| 30S ribosomal protein S13 | 410 | 120 | 3.4 |
| Chaperone protein DnaK | 13 | 4 | 3.3 |
| Enolase | 3 | 1 | 3.0 |
| Tetratricopeptide repeat protein | 49 | 24 | 2.0 |

**Supplementary Table 2: Proteins with 2-fold higher in strain ATCC 25611 than in strain H31226 by CE/LC MASS.**
